# Noradrenergic Depletion by DSP-4 Reduces Morphological Complexity of Hippocampal Astrocytes

**DOI:** 10.64898/2026.07.06.736739

**Authors:** Garima Virmani, Tithi Bhowmick, Swananda Marathe

## Abstract

**Background:** Norepinephrine (NE) released from locus coeruleus (LC) projections regulates astrocyte structure and function through adrenergic receptor signaling. We previously showed that increasing noradrenergic tone with the NE reuptake inhibitor desipramine increases astrocyte ramification in the molecular layer of the dentate gyrus. However, whether tonic LC-derived noradrenergic tone is required to maintain astrocyte morphological complexity *in vivo*, and whether β-adrenergic receptor activation is the effector pathway, remained unclear.

**Methods:** Adult male C57BL/6J mice received DSP-4 (50 mg/kg × 3 days i.p.), a selective LC neurotoxin, with or without concurrent isoproterenol that continued for 21 additional days post cessation of DSP-4 treatment (ISO; 2 mg/kg/day × 24 days), or saline (n = 4 mice per group). Animals were sacrificed 22 days after the final DSP-4 injection. Noradrenergic denervation was confirmed by dopamine β-hydroxylase (DBH) immunostaining. GFAP-immunostained astrocytes in the molecular layer of the dentate gyrus were morphologically characterized using Sholl analysis. Astrocyte density was quantified by SOX9 immunostaining.

**Results:** DSP-4 produced >83% reduction in DBH fiber coverage in the molecular layer. Sholl analysis revealed significant reductions in astrocyte branching complexity in both treatment groups, with the reductions concentrated at distances of 5–15 µm from the soma. The maximum number of intersections was also significantly reduced in both groups. Unexpectedly, ISO did not rescue morphological complexity. While DSP-4 alone did not alter astrocyte density, as measured by the number of SOX9-expressing astrocytes, DSP-4+ISO increased SOX9-positive cell density, dissociating the effects of adrenergic signaling on morphology from those on cell numbers.

**Conclusions:** LC-derived noradrenergic tone is required for the maintenance of astrocyte arbour complexity in the dentate gyrus molecular layer. β-adrenergic receptor activation alone is insufficient to restore structural integrity following noradrenergic denervation, yet promotes astrocyte density independently of structural remodeling. These findings have implications for understanding how LC neurodegeneration in Alzheimer’s disease and depression may compromise hippocampal astrocyte structure and function.

## Introduction

Astrocytes are morphologically complex cells whose elaborate processes tile the hippocampal neuropil in non-overlapping territorial domains, ensheathing synapses and forming the tripartite synapse structure essential for glutamate clearance, potassium buffering, and synaptic modulation (1,2). The structural complexity of hippocampal astrocytes is not static — it undergoes activity-dependent and neuromodulator-driven remodeling that has been implicated in sleep, the pathophysiology of major depressive disorder (MDD), the mechanism of antidepressant action, and several other disease phenotypes (3,4).

Among the neuromodulatory systems regulating astrocyte morphology, norepinephrine (NE) released from locus coeruleus (LC) projections is a prominent candidate. Astrocytes express both α- and β-adrenergic receptors, and noradrenergic signaling regulates GFAP expression, astrocyte volume, and process dynamics (5–7). Activation of β-adrenergic receptors promotes astrocyte process extension in culture (7). Specifically in the hippocampus, we have demonstrated that chronic treatment with the NE reuptake inhibitor desipramine (DMI) significantly increases astrocyte ramification in the molecular layer of the dentate gyrus — the subfield receiving dense LC-derived noradrenergic innervation — and that chronic stress reduces astrocyte complexity in this same region (4,8).

Given that the astrocytes from the molecular layer of the hippocampal dentate gyrus were found to be most vulnerable to the effects of stress as well as desipramine treatment (4,8), we asked whether tonic noradrenergic tone from the LC is required to maintain astrocyte morphological integrity in the molecular layer *in vivo*. LC neurodegeneration is an early and prominent feature of Alzheimer’s disease and is increasingly recognized as a component of MDD pathophysiology (9). Understanding whether LC-derived NE supports astrocyte structure is directly relevant to how noradrenergic depletion in these conditions may compromise hippocampal circuit function through glial remodeling.

DSP-4 (N-(2-chloroethyl)-N-ethyl-2-bromobenzylamine) is a selective neurotoxin that depletes noradrenergic terminals originating from the LC by alkylating the NE reuptake transporter, producing a long-lasting reduction in terminal NE release in LC-innervated brain regions (10). We used DSP-4 to examine the effects of noradrenergic denervation on astrocyte morphology in the molecular layer of the dentate gyrus, and concurrently tested whether sustained β-adrenergic receptor stimulation with isoproterenol (ISO) could rescue any observed morphological deficits.

## Methods

### Animals

Adult male C57BL/6J mice (25–32 g, 6–8 months) were group-housed under a 12h light/dark cycle with *ad libitum* access to food and water. All procedures were approved by the Institutional Animal Ethics Committee (IAEC) of the Indian Institute of Science, Bangalore (CAF/Ethics/738/2019). Four mice were used per experimental group.

### Experimental Design and Drug Treatments

Mice were assigned to three groups: namely Saline (SAL), DSP-4 and DSP-4 + isoproterenol (ISO). DSP-4 and DSP-4 + ISO groups received DSP-4 (Sigma-Aldrich, C8417; 50 mg/kg, i.p.), one injection per day for three consecutive days (days 1–3). The SAL group received an equivalent volume of 0.9% saline solution. 30 minutes after the DSP-4 injection, DSP-4 + ISO group received isoproterenol (Sigma-Aldrich, I6504; 2 mg/kg, i.p.) once daily, while SAL and DSP-4 groups received an equivalent volume of 0.9% Saline solution. Isoproterenol and Saline injections were continued for 21 additional days after DSP-4 treatment ended. 24 hours after the final isoproterenol or saline injections (on day 25), the mice were sacrificed by transcardial perfusion with 0.9% saline, followed by 4% paraformaldehyde (PFA). Brains were post-fixed overnight at 4°C in 4% PFA, cryoprotected in 30% sucrose in 0.1 M phosphate buffer, and sectioned coronally at 40 µm on a sliding microtome (Leica SM2010R).

### Immunohistochemistry

Free-floating sections were blocked in 3% BSA / 1% normal donkey serum / 0.3% Triton X-100 in 0.1M phosphate buffer for 1h at room temperature. For GFAP staining, sections were incubated overnight at 4°C with chicken polyclonal anti-GFAP (1:1000; Novus, NBP1-05198) followed by goat anti-chicken IgY Alexa Fluor 594 (1:1000; Abcam, ab150172) for 2h at room temperature. For SOX9 staining, goat polyclonal anti-SOX9 (1:500; R&D Systems, AF3075) was used with donkey anti-goat IgG Alexa Fluor 568 secondary (1:500; Abcam, ab175474). For DBH staining, Rabbit polyclonal anti-DBH antibody (1:500; Abcam, ab96615) was used with Donkey anti-rabbit Alexa Fluor 488 secondary (1:500; Abcam, ab150073). All sections were counterstained with DAPI and mounted on glass slides with Fluoromount-G.

### Confocal imaging

GFAP-stained sections were imaged on a Zeiss LSM 880 Airyscan confocal microscope with a Plan-Apochromat 20x/0.8 M27 objective. 1024 × 1024 pixels, 8-bit images were acquired at 2µm z plane intervals. DBH- and SOX9-stained sections were imaged on the same system at 20× magnification. The region of interest was defined as the molecular layer above the upper blade of the dentate gyrus granule cell layer.

### Astrocyte morphological analysis

Individual GFAP-immunopositive astrocytes were manually cropped from confocal z-stacks in x, y, and z dimensions using ImageJ (NIH). Maximum intensity projections (MIPs) were generated and processed using PureDenoise (ImageJ plugin), Gaussian blur (σ = 0.5), and Li thresholding. Binary images were skeletonized using the Skeletonize plugin with hole-filling. Skeletonized images were analyzed using SMorph software (4) to extract Sholl intersection profiles and morphometric parameters. All image sets were analyzed blind to the treatment group.

Final group sizes after exclusion of one statistical outlier (identified by the ROUT method, Q = 1%) from the DSP-4 group were: SAL n = 43 cells, DSP-4 n = 36 cells, DSP-4+ISO n = 31 cells, sampled evenly from 4 mice per group. The outlier was confirmed using Gaussian mixture modeling of the principal component 1 (PC1) scores for the full Sholl vector.

### DBH fiber quantification

MIPs of DBH-stained sections were used to quantify the percentage area coverage of DBH-positive fibers within manually drawn ROIs in the molecular layer. Area coverage was calculated using ImageJ thresholding and particle analysis.

### Astrocyte density

SOX9-positive astrocytes were counted within manually defined ROIs in the molecular layer from MIPs of SOX9-stained sections. Counts were divided by ROI area (mm^2^). n = 4 mice per group.

### Statistical analysis

All analyses were performed in Python (SciPy 1.17; scikit-learn 1.8). Data are expressed as mean ± SEM. The overall Sholl profile was compared between groups using multivariate permutation testing on the full Sholl vector (5,000 permutations). To localize the spatial distribution of differences, per-radius Mann-Whitney U tests (two-tailed) were conducted and reported descriptively. Derived morphometric parameters were compared by the Kruskal-Wallis test followed by pairwise Mann-Whitney U tests (two-tailed). DBH fiber coverage and SOX9 density were compared using one-way ANOVA, followed by Dunnett’s multiple comparisons test. Statistical significance threshold was p < 0.05 throughout.

## Results

### DSP-4 depletes noradrenergic terminals in the molecular layer of the dentate gyrus

We treated chronically mice with DSP-4 and ISO as explained in the methods section (Figure 1A). To confirm effective noradrenergic denervation, we quantified DBH-immunopositive fiber area coverage in the molecular layer (Figure 1B). In the molecular layer, both DSP-4 and DSP-4+ISO produced a statistically significant and dramatic reduction in DBH fiber coverage (SAL: 11.59 ± 0.38%, DSP-4: 1.96 ± 0.45%, DSP-4+ISO: 2.18 ± 0.44%; both p < 0.0001 vs SAL; Figure 1C). The near-complete loss of DBH fibers in the molecular layer (>83% reduction) confirms effective LC-derived noradrenergic denervation. These data establish that DSP-4 treatment caused significant noradrenergic denervation in the molecular layer of the hippocampal dentate gyrus.

**Figure 1.**
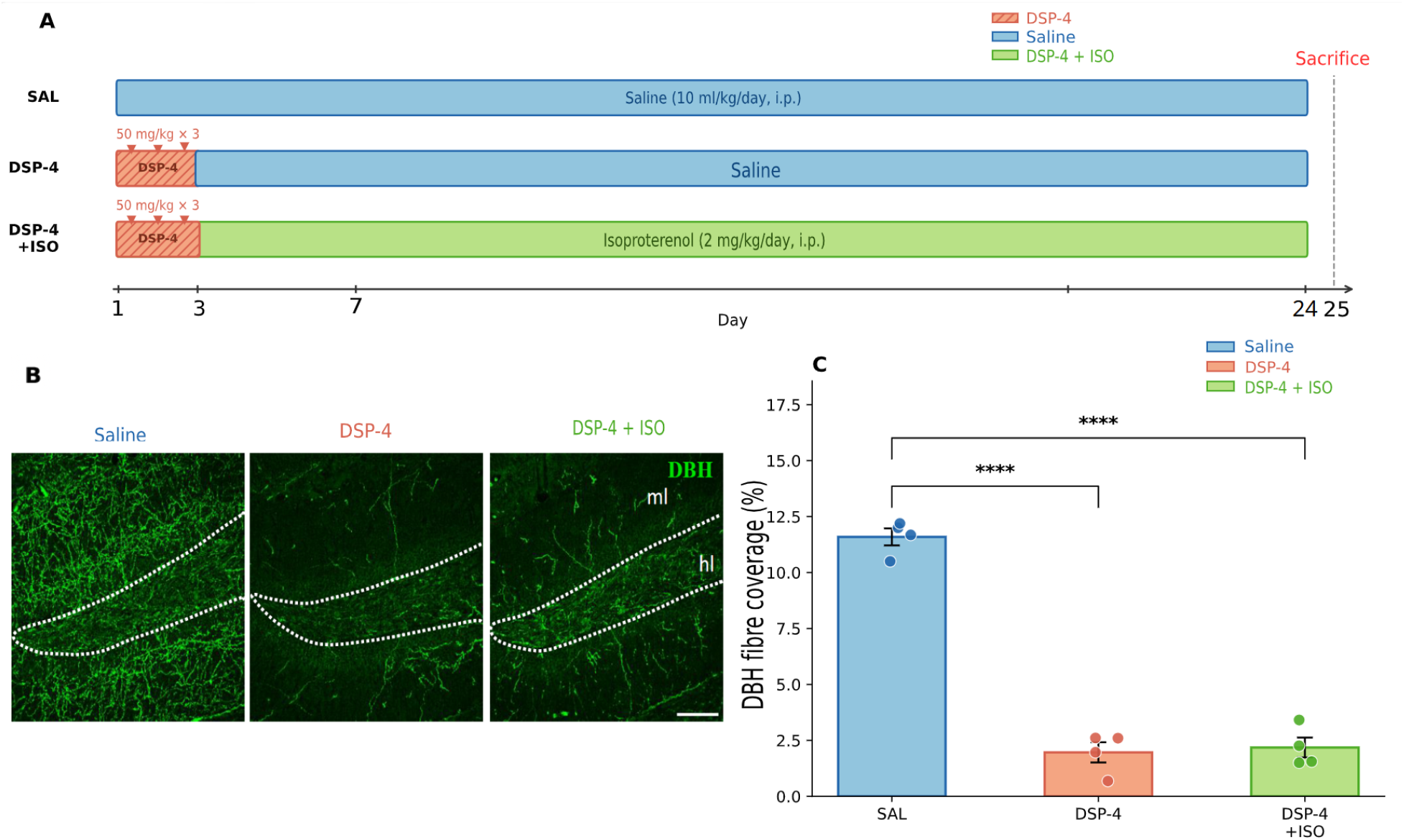
DSP-4 selectively depletes noradrenergic terminals in the molecular layer of the dentate gyrus. (A) Schematic of the treatment paradigm. Three groups were used: Saline (SAL), DSP-4, and DSP-4+isoproterenol (DSP-4+ISO). DSP-4 (50 mg/kg, i.p.) was administered on days 1–3 in both treatment groups. The DSP-4+ISO group received concurrent isoproterenol (ISO; 2 mg/kg/day, i.p.) from day 1 through day 24. The SAL group received saline (10 ml/kg/day, i.p.) for 24 days. All animals were sacrificed on day 25. Downward triangles indicate DSP-4 injection days. (B) Representative confocal images of dopamine β-hydroxylase (DBH) immunostaining in the molecular layer of the dentate gyrus in SAL, DSP-4, and DSP-4+ISO groups. Scale bar: 100 µm. (C) Quantification of DBH-positive fibre area coverage (%) in the molecular layer. Both DSP-4 and DSP-4+ISO groups show a dramatic and significant reduction in DBH fibre coverage compared to saline controls (SAL: 11.59 ± 0.38%, DSP-4: 1.96 ± 0.45%, DSP-4+ISO: 2.18 ± 0.44%; SAL vs DSP-4: p < 0.0001; SAL vs DSP-4+ISO: p < 0.0001; one-way ANOVA F(2,9) = 166.1, p < 0.0001, Dunnett’s post-hoc test). Bars represent mean ± SEM; scatter points show individual mice. n = 4 mice per group. ****p < 0.0001.

### DSP-4 reduces astrocyte Sholl profile complexity

To examine whether noradrenergic denervation affects astrocyte morphological complexity, we performed Sholl analysis on GFAP-immunostained astrocytes in the molecular layer 22 days after final DSP-4 administration (Figure 1A). Both DSP-4 and DSP-4+ISO groups showed consistent reductions in Sholl intersection counts across the profile compared to saline controls. Per-radius Mann-Whitney U tests identified significant differences between SAL and DSP-4 at radii 5, 6, 7, 9, 10, and 13 µm, and between SAL and DSP-4+ISO at radii 5, 6, 7, 8, 9, 10, and 12 µm (all p < 0.05; Figure 2A). The significant radii cluster within 5–15 µm from the soma, corresponding to the proximal-to-mid arbor zone where perisynaptic astrocyte process density is greatest. Whole-curve multivariate permutation testing confirmed the overall Sholl profile differed significantly between groups (SAL vs DSP-4: p = 0.006; SAL vs DSP-4+ISO: p = 0.001; 5,000 permutations).

**Figure 2.**
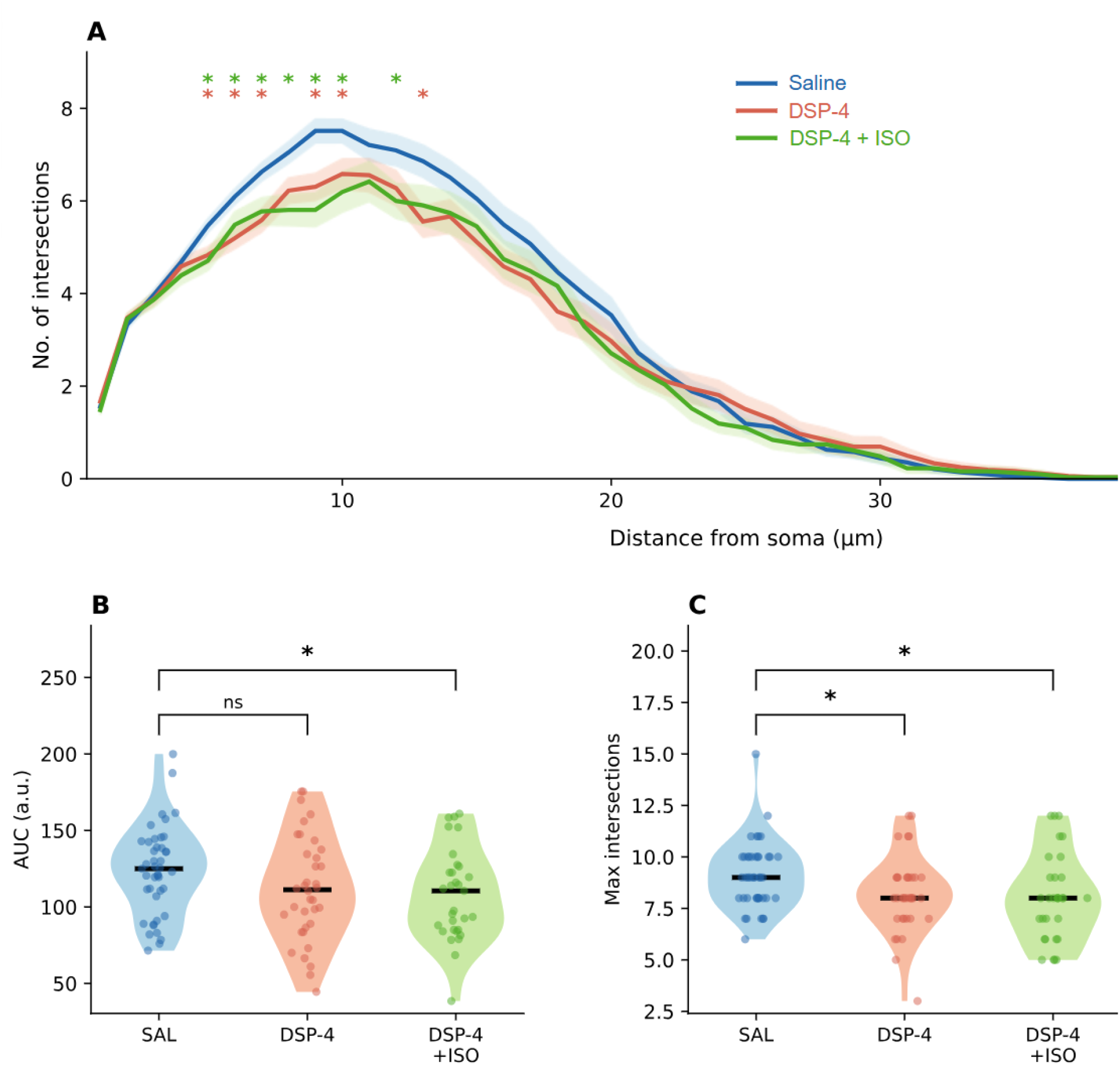
DSP-4 reduces astrocyte morphological complexity in the molecular layer of the dentate gyrus. (A) Mean Sholl profiles (± SEM) for SAL (n = 43 cells), DSP-4 (n = 36 cells), and DSP-4+ISO (n = 31 cells) groups. Coloured asterisks above the profile indicate radii at which intersection counts differed significantly from SAL by per-radius Mann-Whitney U test (two-tailed, p < 0.05): red asterisks for DSP-4 (radii 5, 6, 7, 9, 10, 13 µm), green for DSP-4+ISO (radii 5, 6, 7, 8, 9, 10, 12 µm). The overall Sholl curve differed significantly from SAL by multivariate permutation test on the full Sholl vector (SAL vs DSP-4: p = 0.006; SAL vs DSP-4+ISO: p = 0.001; 5,000 permutations). (B) Area under the Sholl curve (AUC). SAL vs DSP-4+ISO: p = 0.035, Cohen’s d = 0.53; SAL vs DSP-4: p = 0.127, Cohen’s d = 0.37 (Mann-Whitney U, two-tailed). (C) Maximum intersections. SAL vs DSP-4: p = 0.020, Cohen’s d = 0.53; SAL vs DSP-4+ISO: p = 0.016, Cohen’s d = 0.56 (Mann-Whitney U, two-tailed). Kruskal-Wallis: AUC H = 4.80, p = 0.09; maximum intersections H = 8.12, p = 0.017. Violin plots show the full distribution of individual cell values; horizontal lines indicate group medians; scatter points show individual cells. Italic d values indicate Cohen’s d effect size relative to Saline. *p < 0.05; ns = not significant. Cells were sampled evenly from 4 mice per group.

### DSP-4 reduces astrocyte maximum intersections and area under the Sholl curve

Derived Sholl parameters quantified the magnitude of the morphological simplification (Figures 2B–C). Area under the Sholl curve (AUC) showed a significant reduction in the DSP-4+ISO group (SAL: 123.5 ± 3.9 a.u., DSP-4: 111.8 ± 5.0 a.u., DSP-4+ISO: 107.9 ± 4.6 a.u.; SAL vs DSP-4+ISO p = 0.035, Cohen’s d = 0.53; SAL vs DSP-4 p = 0.127, Cohen’s d = 0.37; Figure 2B). Maximum intersections were significantly reduced in both treatment groups compared to saline (Kruskal-Wallis H = 8.12, p = 0.017; SAL: 9.09 ± 0.22, DSP-4: 8.17 ± 0.27, DSP-4+ISO: 8.03 ± 0.33; Mann-Whitney U: SAL vs DSP-4 p = 0.020, Cohen’s d = 0.53; SAL vs DSP-4+ISO p = 0.016, Cohen’s d = 0.56; Figure 2C). The medium effect sizes (Cohen’s d 0.37–0.56) across both parameters are consistent with a genuine but graded reduction in astrocyte complexity rather than catastrophic structural collapse.

These results indicate that sustained activation of β-adrenergic receptors is insufficient to prevent or reverse the structural consequences of noradrenergic denervation in the molecular layer.

### DSP-4 does not alter astrocyte number, but DSP-4+ISO increases astrocyte density in the molecular layer

To determine whether the morphological changes reflected structural remodeling of individual cells or changes in astrocyte population size, we quantified the density of SOX9-positive astrocytes in the molecular layer (Figure 3A). DSP-4 alone did not alter astrocyte density (SAL: 1,189 ± 55, DSP-4: 1,269 ± 87 cells/mm^2^; p = 0.595; Figure 3B). However, the DSP-4+ISO group showed an increase in SOX9-positive cell density compared to saline controls (1,432 ± 39 cells/mm^2^; p = 0.044). The absence of a change in astrocyte number with DSP-4 alone confirms that the morphological simplification we observe reflects structural remodeling of individual cells rather than cell loss. The selective increase in astrocyte density with DSP-4+ISO, dissociated from any structural rescue, suggests that ISO engages astrocyte proliferative or survival pathways independently of the cytoskeletal remodeling pathway that maintains process complexity.

**Figure 3.**
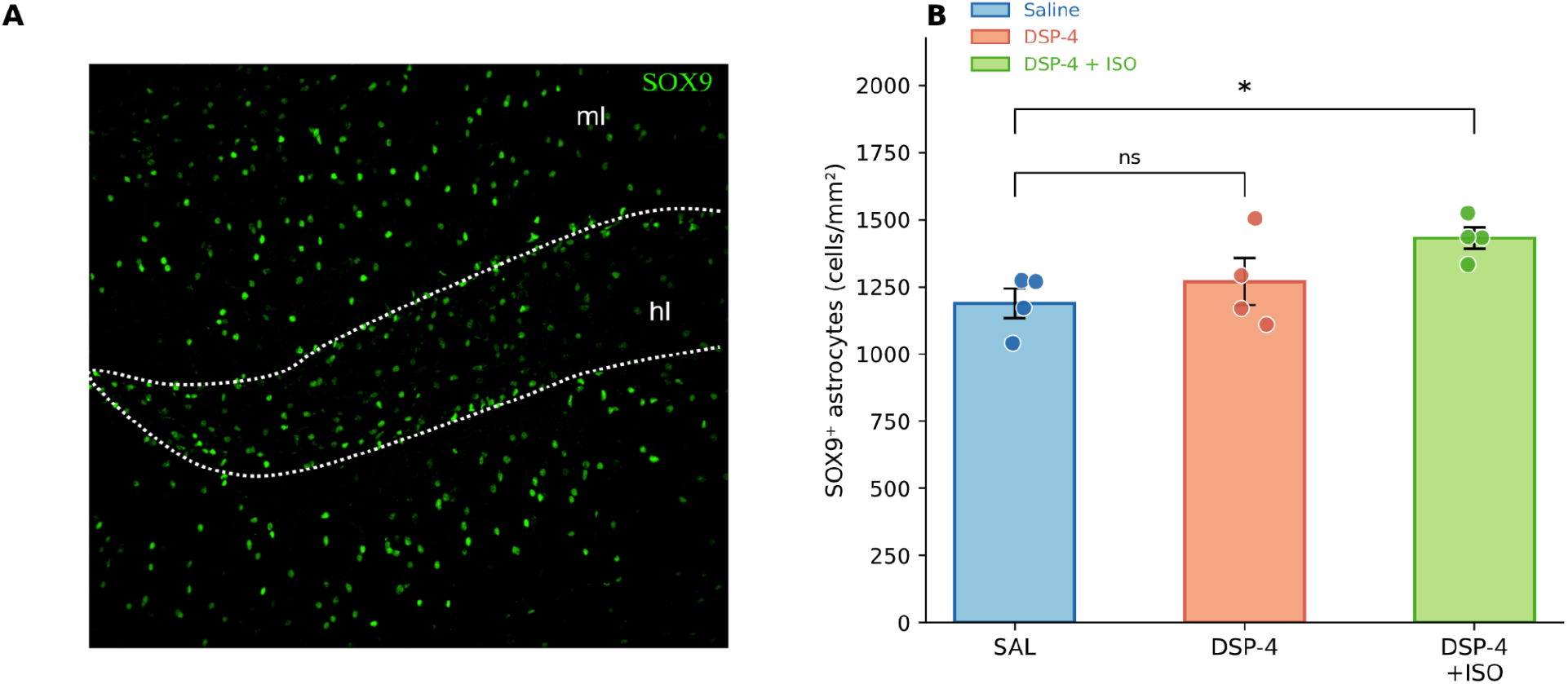
DSP-4 does not alter astrocyte number; DSP-4+ISO increases SOX9^+^ astrocyte density in the molecular layer. (A) Representative confocal image of SOX9 immunostaining in the molecular layer of the dentate gyrus. Scale bar: 100 µm. (B) Quantification of SOX9-positive astrocyte density (cells/mm^2^) in the molecular layer. DSP-4 alone did not significantly alter astrocyte density compared to saline controls (SAL: 1,189 ± 55, DSP-4: 1,269 ± 87 cells/mm^2^; p = 0.595). DSP-4+ISO significantly increased SOX9-positive cell density relative to saline (1,432 ± 39 cells/mm^2^; p = 0.044). Bars represent mean ± SEM; scatter points show individual mice. n = 4 mice per group. One-way ANOVA F(2,9) = 3.77, p = 0.065; Dunnett’s post-hoc test. *p < 0.05; ns = not significant.

## Discussion

We demonstrate that selective depletion of LC-derived noradrenergic terminals by DSP-4 significantly reduces the morphological complexity of GFAP-positive astrocytes in the molecular layer of the mouse dentate gyrus, as assessed 22 days after treatment. The effect is concentrated at proximal-to-mid arbor radii (5–15 µm from the soma), consistent with selective retraction of perisynaptic astrocyte processes. Critically, this morphological simplification occurs without loss of astrocytes — cell density is unchanged by DSP-4 alone — indicating genuine structural remodeling of individual cells rather than a population-level change.

The molecular layer of the dentate gyrus shows >83% reduction in DBH fiber coverage (p < 0.0001). This anatomical correspondence is mechanistically coherent: astrocytes in the molecular layer are exposed to substantially higher NE levels from LC terminals under normal conditions and are therefore more severely affected by its withdrawal. This subfield specificity also aligns with our previous findings that the molecular layer is the dentate subfield most responsive to noradrenergic pharmacological manipulation — chronic DMI treatment increases astrocyte ramification predominantly in this region (4) — suggesting a bidirectional relationship between noradrenergic tone and astrocyte structural complexity in this zone.

The proximal-to-mid arbor localization of the effect (5–15 µm from soma) is biologically significant. This zone corresponds to the primary perisynaptic astrocyte process (PAP) compartment, where glutamate transporters GLT-1 and GLAST are concentrated, and the tripartite synapse interface is established (2). Process retraction at these radii is expected to reduce the astrocyte’s capacity for glutamate uptake and for potassium spatial buffering at perforant path synapses in the molecular layer. While we have not directly measured glutamate transporter expression or synaptic transmission in this study, the structural changes we report provide a morphological basis for predicting functional consequences at the perforant path–dentate synapses following LC denervation. This prediction is testable in future work using electrophysiological or biochemical approaches.

The failure of isoproterenol to rescue the DSP-4-induced morphological simplification is a substantive negative finding. ISO is a non-selective β-adrenergic agonist that bypasses the NE reuptake step blocked by DSP-4, providing direct receptor stimulation at both β1- and β2-adrenergic receptors on astrocytes. Its failure to prevent or reverse morphological simplification argues against a simple model in which β-adrenergic signaling is the sole effector of NE-dependent astrocyte structural maintenance. Several mechanistic explanations merit consideration. First, noradrenergic terminal loss may reduce co-release of other signaling molecules — including ATP, neuropeptide Y, and galanin — that act synergistically with NE to maintain astrocyte structure (11). Second, tonic versus phasic receptor activation may matter: endogenous NE release is phasic and activity-dependent, while ISO in the current paradigm was administered as a single daily i.p. injection, which cannot replicate the spatiotemporal dynamics of synaptic NE release. Third, α-adrenergic receptors, which mediate opposing effects on cytoskeletal dynamics in astrocytes compared to β receptors (7), are not engaged by ISO, and their contributions to structural maintenance may be independent and non-redundant.

The finding that DSP-4+ISO significantly increases astrocyte density in the molecular layer (p = 0.044) while failing to rescue morphological complexity reveals a dissociation between proliferative/survival effects and structural remodeling effects of adrenergic signaling on astrocytes. β-adrenergic receptor activation is known to promote astrocyte survival and proliferation through cAMP-PKA pathways (12), and our data suggest these population-level effects can occur independently of the cytoskeletal remodeling required for process elaboration. This dissociation has implications for therapeutic strategies targeting astrocyte function in the context of LC degeneration: increasing astrocyte numbers may not compensate for the functional deficit resulting from the morphological simplification of individual cells.

A limitation of the present study is the use of cell-level statistics with n = 4 animals per group, which limits power for animal-level analyses. The effects are consistent in direction across individual animals, and the whole-curve permutation test confirms the Sholl profile difference is unlikely to arise by chance (p = 0.001–0.006). Effect sizes at the cell level are medium (Cohen’s d 0.37–0.56), consistent with a genuine but graded structural change. Confirmation in larger cohorts and extension to female mice would strengthen the generalisability of these findings. Additionally, GFAP-based Sholl analysis captures only the core cytoskeletal scaffold (∼15% of astrocyte volume) and cannot resolve the finest perisynaptic processes; their assessment would require genetic membrane-labeling approaches.

In summary, tonic LC-derived noradrenergic tone is required for the maintenance of astrocyte morphological complexity in the hippocampal molecular layer, and this structural role cannot be substituted by exogenous β-adrenergic receptor activation alone. These findings contribute to understanding how LC neurodegeneration — a prominent early feature of Alzheimer’s disease and a component of the pathophysiology of depression — may compromise hippocampal astrocyte structure and, by inference, synaptic function in the molecular layer of the dentate gyrus.

